# Unbiased estimation of linkage disequilibrium from unphased data

**DOI:** 10.1101/557488

**Authors:** Aaron P. Ragsdale, Simon Gravel

## Abstract

Linkage disequilibrium is used to infer evolutionary history, to identify genomic regions under selection, and to dissect the relationship between genotype and phenotype. In each case, we require accurate estimates of linkage disequilibrium statistics from sequencing data. Unphased data present a challenge because multi-locus haplotypes cannot be inferred exactly. Widely used estimators for the common statistics *r*^2^ and *D*^2^ exhibit large and variable upward biases that complicate interpretation and comparison across cohorts. Here, we show how to find unbiased estimators for a wide range of two-locus statistics, including *D*^2^, for both single and multiple randomly mating populations. These unbiased statistics are particularly well suited to estimate effective population sizes from unlinked loci in small populations. We develop a simple inference pipeline and use it to refine estimates of recent effective population sizes of the threatened Channel Island Fox populations.

## Introduction

Linkage disequilibrium (LD), the association of alleles between two loci, is informative about evolutionary and biological processes. Patterns of LD are used to infer past demographic events, identify regions under selection, estimate the landscape of recombination across the genome, and discover genes associated with biomedical and phenotypic traits. These analyses require accurate and efficient estimation of LD statistics from genome sequencing data.

LD is typically given as the covariance or correlation of alleles between pairs of loci. Estimating this covariance from data is simplest when we directly observe haplotypes (in haploid or phased diploid sequencing), in which case we know which alleles co-occur on the same haplotype. However, most whole-genome sequencing of diploids is unphased, leading to ambiguity about the co-segregation of alleles at each locus.

The statistical foundation for computing LD statistics from unphased data that was developed in the 1970s (Hill, 1974; Cockerham and Weir, 1977; Weir, 1979) has led to widely used approaches for their estimation from modern sequencing data (Excoffier and Slatkin, 1995; Rogers and Huff, 2009). While these methods provide accurate estimates for the covariance and correlation (*D* and *r*), they do not extend to other two-locus statistics, and they result in biased estimates of *r*^2^ (Waples, 2006). This bias confounds interpretation of *r*^2^ decay curves.

Here, we extend an approach for estimating the covariance *D* introduced by Weir (1979) to find unbiased estimators for a large set of two-locus statistics including *D*^2^ and 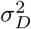. We show that these estimators are accurate for the low-order LD statistics used in demographic and evolutionary inferences. We provide an estimator for *r*^2^ with improved qualitative and quantitative behavior over the widely used approach of Rogers and Huff (2009), although it remains a biased estimator. In general, for analyses sensitive to biases in the estimates of statistics, we recommend the use of *D*^2^ or 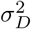 over *r*^2^.

As a concrete use case, we consider estimating recent effective population size (*N*_*e*_) from observed LD between unlinked loci, a common analysis when population sizes are small, typical in conservation and domestication genomics studies. Waples (2006) suggested combining an empirical bias correction for estimates of *r*^2^ with an approximate theoretical result from Weir and Hill (1980) to estimate *N*_*e*_.

Here, we propose an alternative approach to estimate *N*_*e*_ using our unbiased estimator for 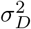 that avoids many of the assumptions and biases associated with *r*^2^ estimation. We first derive expectations for 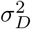 and related statistics between unlinked loci and compare estimates of *N*_*e*_ based on 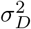 and *r*^2^ using simulated data. As an application, we reanalyze sequencing data from Funk et al. (2016) to estimate recent *N*_*e*_ in the threatened Channel Island fox populations using 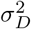. Our estimates are overall consistent with those reported in Funk et al. (2016) using the approach from (Waples, 2006), with the exception of the San Nicolas Island population where the 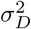-based estimate of 13.8 individuals is over 6 times larger and 100 standard deviations away from the *r*^2^-based estimate of 2.1. Our analysis further suggests population structure or recent gene flow into island fox populations.

## Linkage disequilibrium statistics

For measuring LD between two loci, we assume that each locus carries two alleles: *A/a* at the left locus and *B/b* at the right locus. We think of *A* and *B* as the derived alleles, although the expectations of statistics that we consider are unchanged if alleles are randomly labeled instead. Allele *A* has frequency *p* in the population (allele *a* has frequency 1 − *p*), and *B* has frequency *q* (*b* has 1 − *q*). There are four possible two-locus haplotypes, *AB*, *Ab*, *aB*, and *ab*, whose frequencies sum to 1.

For two loci, LD is typically given by the covariance or correlation of alleles co-occurring on a haplotype. The covariance is denoted *D*:

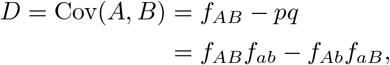

and the correlation is denoted *r*:

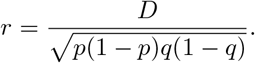

Squared covariances (*D*^2^) and correlations (*r*^2^) see wide use in genome-wide association studies to thin data for reducing correlation between SNPs and to characterize local levels of LD (e.g. Speed et al. (2012)). While the average of *D* across sites is 0 under broad conditions, averages of *D*^2^ and *r*^2^ across sites are informative about demography: the scale and decay rate of *r*^2^ curves reflect population sizes over a range of time periods (Tenesa et al., 2007; Hollenbeck et al., 2016), while recent admixture will lead to elevated long-range LD (Moorjani et al., 2011; Loh et al., 2013).

To measure the scale and decay rate of LD statistics, we will compute averages over many independent pairs of loci. To build theoretical predictions for these observations, we take expectations 𝔼[*r*^2^] and 𝔼[*D*^2^] over multiple realizations of the evolutionary process.

It is difficult to compute model predictions for *r*^2^, however, even in simple evolutionary scenarios. Here we consider a related measure proposed by Ohta and Kimura (1969),

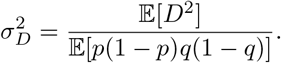

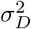 is not as commonly used or reported, although we can compute its expectation from models (Hill and Robertson, 1968) and estimate it from data (as described in this study). Recent studies have demonstrated that 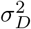 can be used to infer population size history (Rogers, 2014) and, along with a set of related statistics, allows for powerful inference of multi-population demography and archaic introgression (Ragsdale and Gravel, 2019).

## Results

### Estimating LD from data

In the Methods, we present an approach to compute unbiased estimators for a broad set of two-locus statistics, for either phased or unphased data. This includes commonly used statistics, such as *D* and *D*^2^, the additional statistics in the Hill-Robertson system (*D*(1 − 2*p*)(1 − 2*q*) and *p*(1 − *p*)*q*(1 − *q*), which we denote *Dz* and *π*_2_, respectively), and, in general, any statistic that can be expressed as a polynomial in haplotype frequencies (*f*’s) or in terms of *p*, *q*, and *D*. We use this same approach to find unbiased estimators for cross-population LD statistics, which were recently used to infer multi-population demographic history (Ragsdale and Gravel, 2019).

We use our estimators for *D*^2^ and *π*_2_ to propose an estimator for *r*^2^ from unphased data, which we denote 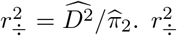 is a biased estimator for *r*^2^ (Discussion). However, it performs favorably in comparison to the common approach of first computing 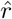 (e.g. via Rogers and Huff (2009)) and simply squaring the result (Figure 1).

**Figure 1:**
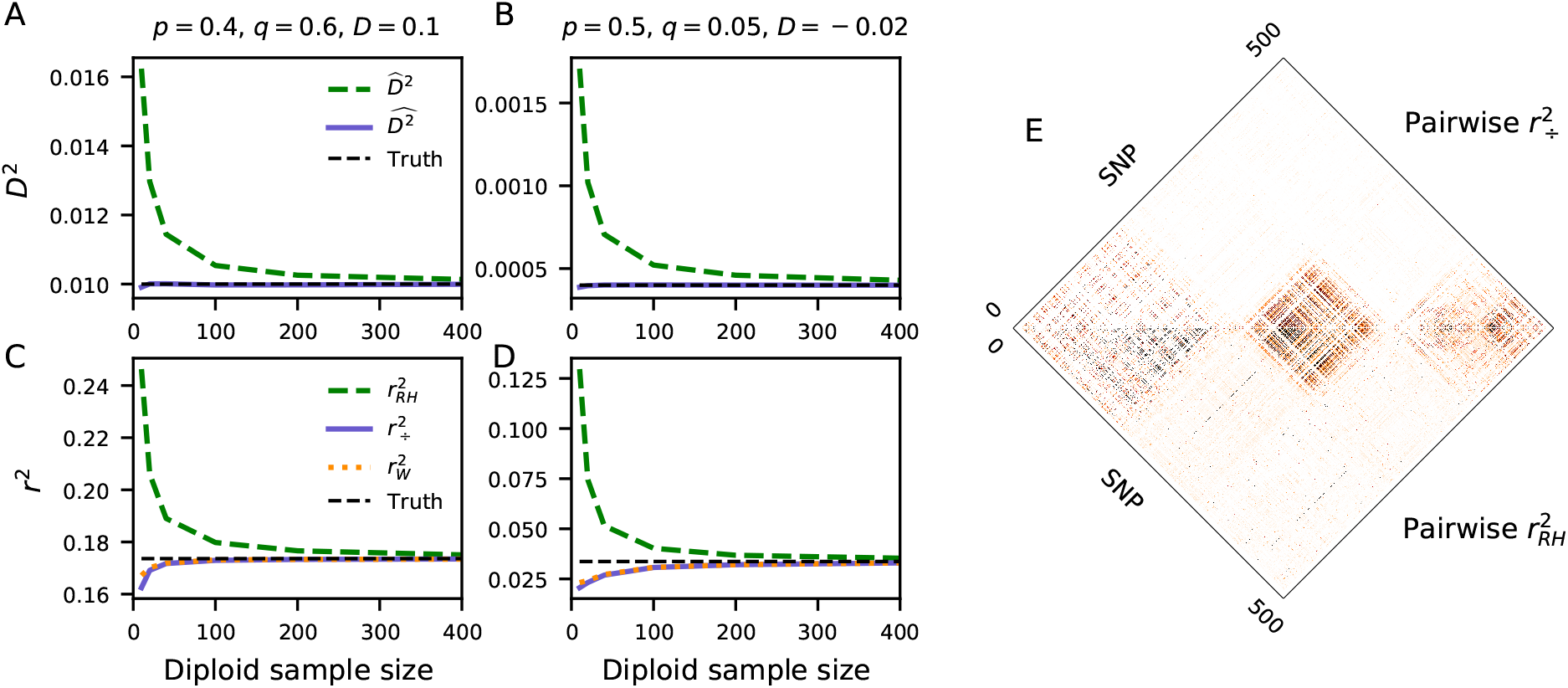
LD estimation. A-B: Computing *D*^2^ by taking the square of the covariance overestimates the true value, while our approach is unbiased for any sample size. C-D: Similarly, computing *r*^2^ by estimating *r* and squaring it (here, via the Rogers-Huff approach, 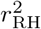) overestimates the true value. Waples proposed empirically estimating the bias due to finite sample size and subtracting this bias from estimated *r*^2^. Our approach, 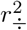 is less biased than 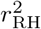 and provides similar estimates to 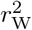 without the need for ad hoc bias correction. (Additional comparisons found in Figure S3.) E: Pairwise comparison of *r*^2^ for 500 neighboring SNPs in chromosome 22 in CHB from 1000 Genomes Project Consortium et al. (2015). 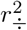 (top) and 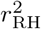 (bottom) are strongly correlated, although 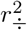 displays less spurious background noise.

To explore the performance of this estimator, we first simulated varying diploid sample sizes with direct multinomial sampling from known haplotype frequencies (Figures 1A-D and S3). Estimates of *D*^2^ were unbiased as expected, and 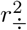 converged to the true *r*^2^ faster than the Rogers-Huff approach. Standard errors of our estimator were nearly indistinguishable from Rogers and Huff (2009) (Figure S1), and the variances of estimators for statistics in the Hill-Robertson system decayed with sample size as 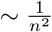 (Figure S2).

Second, we simulated 1 Mb segments of chromosomes under steady state demography (using `msprime` (Kelleher et al., 2016)) to estimate *r*^2^ decay curves using both approaches. Our estimator was invariant to phasing and displayed the proper decay properties in the large recombination distance limit (Figure 2A). With increasing distance between SNPs, 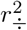 approached zero as expected, while the Rogers-Huff *r*^2^ estimates converged to positive values, as expected in a finite sample (Waples, 2006).

**Figure 2:**
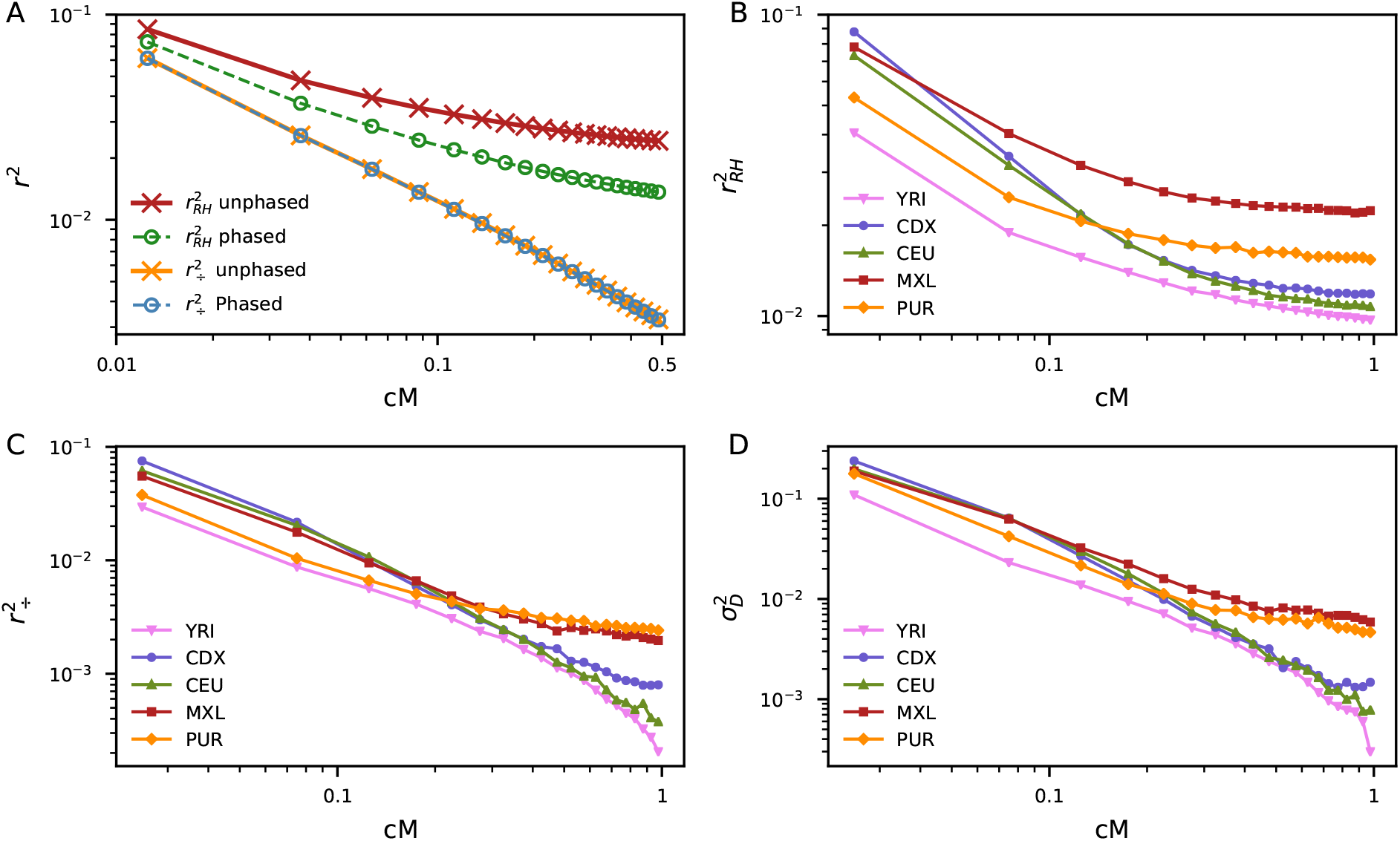
Decay of *r*^2^ with distance. A: Comparison between our estimator 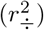 and Rogers and Huff (2009) (RH) under steady state demography. The 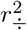-curve displays the appropriate decay behavior and is invariant to phasing, while the RH approach produces upward biased *r*^2^ and is sensitive to phasing. Estimates were computed from 1,000 1Mb replicate simulations with constant mutation and recombination rates (each 2 × 10^−8^ per base per generation) for *n* = 50 sampled diploids using `msprime` (Kelleher et al., 2016). 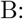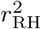 decay for five populations in 1000 Genomes Project Consortium et al. (2015), including two putatively admixed American populations (MXL and PUR), computed from intergenic regions. 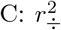 decay for the same populations. D: Decay of 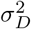 computed using 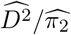. The 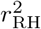 decay curves show excess long-range LD in each population, while our estimator qualitatively differentiates between populations.

Finally, we computed the decay of *r*^2^ across five population from the 1000 Genomes Project Consortium et al. (2015) (Figure 2B-D). 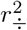 shows distinct qualitative behavior across populations, with recently admixed populations exhibiting distinctive long-range LD. However, *r*^2^ as estimated using the Rogers-Huff approach displayed long-range LD in every population, confounding the signal of admixture in the shape of *r*^2^ decay curves.

### Estimating *N*_*e*_ from LD between unlinked loci

Observed linkage disequilibrium between unlinked markers is widely used to estimate the effective population size (*N*_*e*_) in small populations (Hill, 1981; Waples, 1991, 2006; Waples and Do, 2008; Do et al., 2014). This estimate of *N*_*e*_ reflects the effective number of breeding individuals over the last one to several generations, since LD between unlinked loci is expected to decay rapidly over just a handful of generations. Analytic solutions for 𝔼[*r*^2^] are unavailable, although a classical result uses a ratio of expectations to approximate

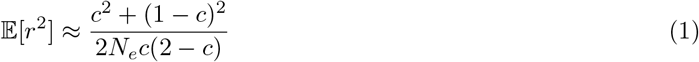

for a randomly mating population, where *c* is the per generation recombination probability between two loci (Equation 3 in Weir and Hill (1980) due to Avery (1978)). For unlinked loci (*c* = 1/2), Equation 1 reduces to 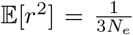 (for a monogamous mating system, 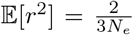 (Weir and Hill, 1980)). Rearranging this equation provides an estimate for *N*_*e*_ if we can estimate *r*^2^ from data.

As pointed out by Waples (2006), failing to account for sample size bias when estimating *r*^2^ from data leads to strong downward biases in 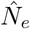. Waples used Burrows’ Δ to estimate 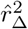 (again following Weir and Hill (1980)) and used simulations to empirically estimate the bias in the estimate due to finite sample size (given by Var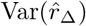). Subtracting this estimated bias from 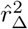 gives an empirically corrected estimate for *r*^2^,

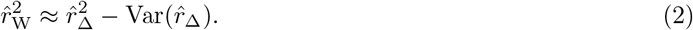

Waples showed that 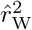 removes much of the bias in *N*_*e*_ estimates (Figure 1C-D). Bulik-Sullivan et al. (2015) used a similar bias correction (via the *δ*-method) that appears to perform comparably to 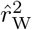 (Figure S3).

Thus, both theoretical predictions for *r*^2^ and its estimation from data are challenging. We show in the next section that these problems disappear if we consider instead the Hill and Robertson (1968) statistic 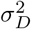.

### Predicting 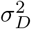 for unlinked and linked loci

The Avery equation (1) was derived under the assumption that the expectation of ratios equals the ratio of expectation. In other words, it assumes that 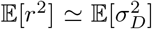. By working directly with 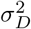, we therefore save both a theoretical approximation and the need for empirical finite sample bias correction. In a random-mating diploid Wright-Fisher model with *c* = 1/2, we show in the Appendix that 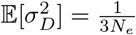, as suggested by the Avery equation, whereas monogamy leads to 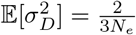. A similar approach allows us to show that 𝔼[*D*(1 − 2*p*)(1 − 2*q*)], another statistic from the Hill-Robertson system, is approximately zero for unlinked loci (its leading-order term is of order 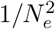).

For tightly linked loci (*c* ≪ 1), Ohta and Kimura (1969) showed that 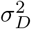 is approximated as

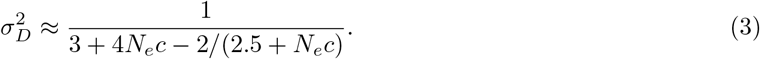

This approximation is accurate for small mutation and recombination rates and for both large and small population sizes at low recombination distances (Figure 3A). Rearranging Equation 3 then provides a direct estimate for *N*_*e*_ for any given recombination distance (Figure 3B), though the approximation is only valid for *c* ≪ 1.

**Figure 3:**
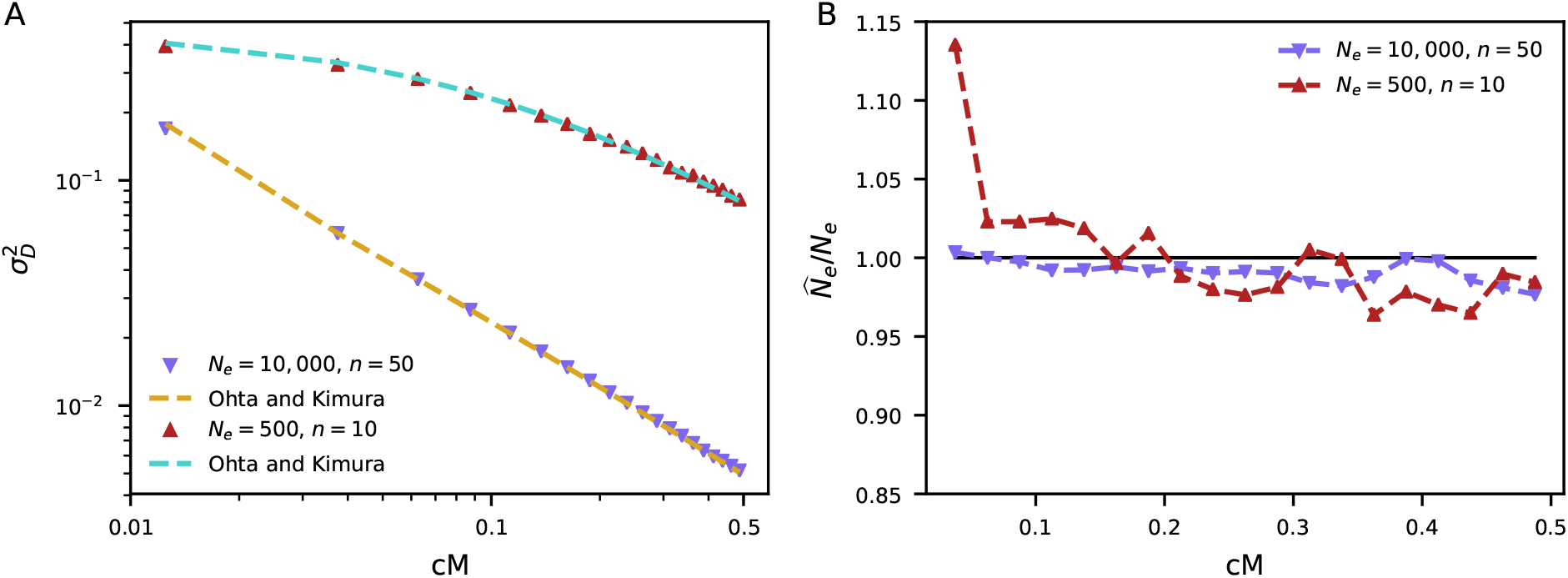
Using 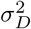 to estimate *N*_*e*_. A: The estimation due to Ohta and Kimura (1969) provides an accurate approximation for 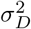 for both large and small sample sizes. Here, we compare to the same simulations used in Figure 2A for *N*_*e*_ =10,000 with sample size *n* = 50 and *N*_*e*_ = 500 with sample size *n* = 10. B: Using 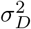 estimated from these same simulations and rearranging Equation 3 provides an estimate for *N*_*e*_ for each recombination bin. The larger variance for *N*_*e*_ = 500 is due to the small sample size leading to noise in estimated 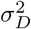.

### Comparison of methods for estimating *N*_*e*_ using simulated data

We simulated data with effective population sizes *N*_*e*_ = 100 and 400 using `fwdpy11` (Thornton, 2014) to compare the performance of inferring 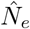 from `NeEstimator` ver. 2.1 (Do et al., 2014), which uses 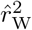, and from 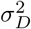 (see Methods for simulation details). Generally, using our estimators for 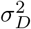 provided less biased estimates of *N*_*e*_ (Figs. 4 and S4). This was the case even when data was filtered by minor allele frequency (MAF), a strategy recommended to reduce bias for `NeEstimator` but that is not required or desirable in the 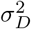 approach. Estimates from 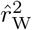 had smaller variance when filtering by MAF, but higher mean squared error for larger sample sizes (Table S1). In practice, `NeEstimator` provides estimates with different cutoff choices and lets the user decide on the best cutoff choice.

**Figure 4:**
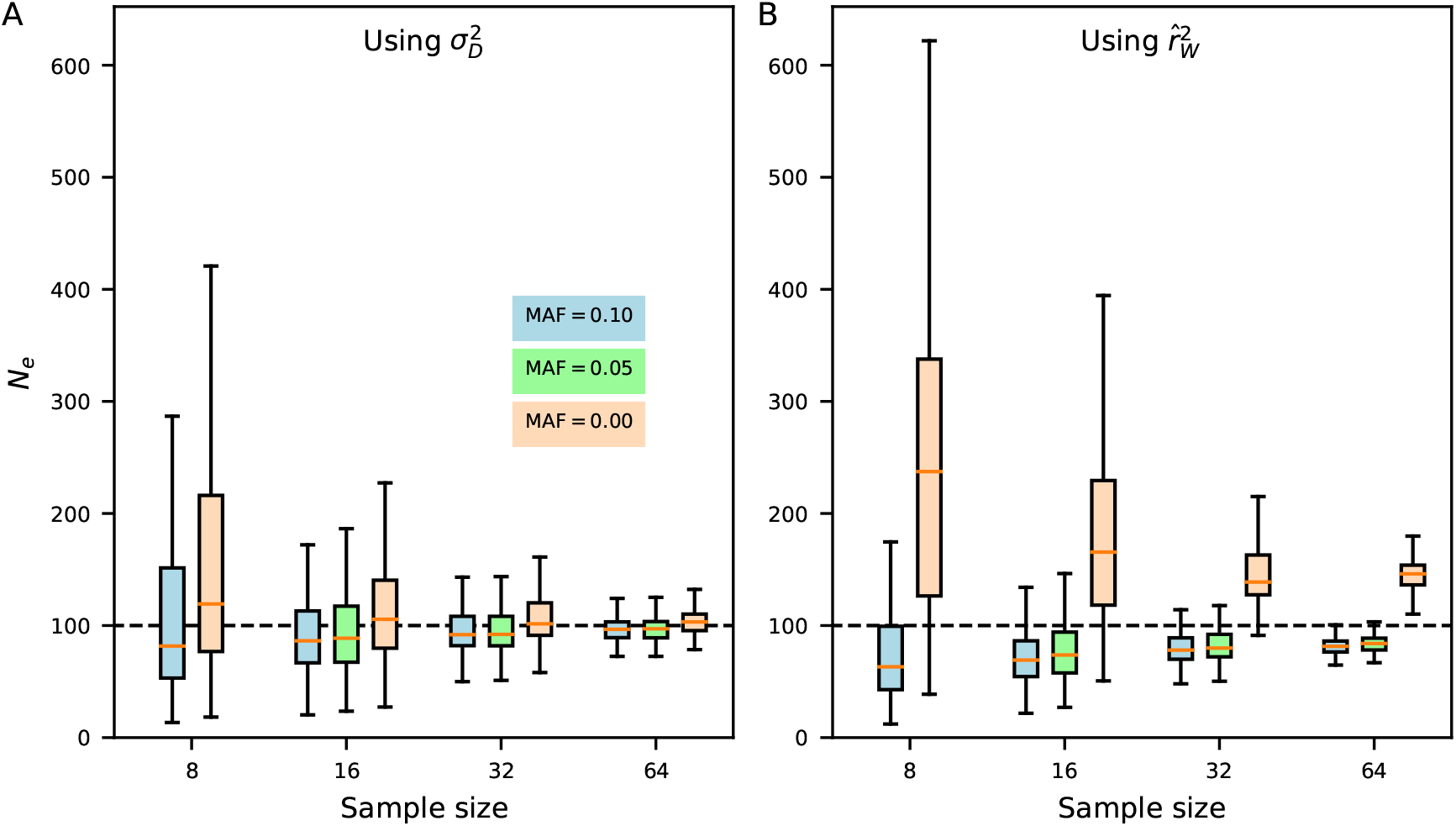
Performance of *N*_*e*_ estimation on simulated data. We used `fwdpy11` (Thornton, 2014) to simulate genotype data for the given sample sizes and *N*_*e*_ = 100 (see Methods for details). Although estimates of *N*_*e*_ using (A) 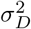 had slightly larger variances than estimates using (B) Eqs. 1 and 2 (computed using `NeEstimator` (Do et al., 2014)), estimates from 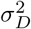 were unbiased when using all data and less biased when filtering by minor allele frequency, resulting in lower MSE (Table S1).

We also explored the effect of inbreeding on estimates of 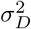 and *σ*_*Dz*_ = 𝔼[*Dz*]/𝔼[*π*_2_] using simulated data. Unsurprisingly, higher rates of inbreeding lead to higher values of 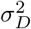 between unlinked loci, which results in deflated estimates of *N*_*e*_ (Figure S5A-B). *σ*_*Dz*_ is robust to inbreeding, with expected value near zero even for large selfing rates (Figure S5C). While *σ*_*Dz*_ cannot be used to provide an estimate for *N*_*e*_ (as its expectation is zero), it could instead be used to distinguish between different violations of model assumptions: if we also measure *σ*_*Dz*_ to be significantly elevated above zero, it might suggest population structure or recent migration into the focal population (Ragsdale and Gravel, 2019).

### The effective population sizes of island foxes

The island foxes (*Urocyon littoralis*) that inhabit the Channel Islands of California have recently experienced severe population declines due to predation and disease. For this reason they have been closely studied to inform protection and management decisions. More generally, they provide an exemplary system to study the genetic diversity and evolutionary history of endangered island populations (Wayne et al., 1991; Coonan et al., 2010; Robinson et al., 2016; Funk et al., 2016; Robinson et al., 2018). A recent study aimed to disentangle the roles of demography (including sharp reductions in population size, resulting in strong genetic drift) and differential selection in shaping the genetics of island foxes across the six Channel Islands (Funk et al., 2016). In addition to genetic analyses based on single-site statistics, Funk et al. (2016) used `NeEstimator` (Do et al., 2014) to infer recent 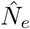 for each of the island fox populations (reproduced in Table 1).

**Table 1:** Inferred island fox effective population sizes. LD between unlinked loci provides an estimate for the effective number of (breeding) individuals in the previous several generations. Funk et al. (2016) used `NeEstimator` (Do et al., 2014) to estimate *N*_*e*_ for six island fox populations in the Channel Islands of California (left). We used this same data to compute *N*_*e*_ using our estimator for 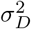 instead (right), obtaining results largely consistent with Funk et al. (2016). Notably, Funk et al. inferred an extremely small size on San Nicolas Island 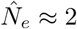, while our estimate is somewhat larger and on the same order of magnitude of 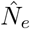 from other islands with small effective population sizes. 90% confidence intervals were computed via 200 resampled bootstrap replicates (Methods).

| Population | $\hat{N}_e$ (95% CI) reported<br>in Funk et al. (2016) | | $\hat{N}_e$ (95% CI) using $\sigma_D^2$ | |
| --- | --- | --- | --- | --- |
| San Miguel I. | 13.7 | (13.2 – 14.1) | 15.3 | (14.5 – 16.1) |
| Santa Rosa I. | 13.6 | (13.5 – 13.7) | 13.3 | (13.0 – 13.6) |
| Santa Cruz I. | 25.1 | (24.6 – 25.5) | 22.8 | (22.4 – 23.3) |
| Santa Catalina I. | 47.0 | (46.7 – 47.4) | 40.9 | (40.4 – 41.6) |
| San Clemente I. | 89.7 | (77.1 – 107.0) | 59.1 | (53.0 – 66.7) |
| San Nicolas I. | 2.1 | (2.0 – 2.2) | 13.8 | (13.0 – 15.2) |

Using the same 5,293 variable sites reported and analyzed in Funk et al. (2016), we computed 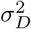 for each of the six island fox populations to estimate *N*_*e*_. Results using 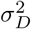 were generally consistent with those computed in Funk et al. (2016) using 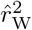 (Tables 1 and S2). Perhaps most notably, the San Nicolas Island population, which was previously inferred to have the extremely small effective size of 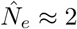, was inferred to have 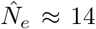. While this is still quite small, it is more similar to the effective sizes inferred in other island fox populations.

We also estimated *σ*_*Dz*_ for each population and found that it was consistently and significantly elevated above zero in each population (Table S2). This suggests that some model assumptions are not being met. From simulated data, neither inbreeding nor filtering by minor allele frequency should result in elevated observed *σ*_*Dz*_ (Figs. S5 and S6). The discrepancy could instead be caused by population substructure or recent migration between populations. It may also be driven by technical artifacts: we analyzed the data with the assumption that the separate RAD contigs were effectively unlinked (reads were not mapped to a reference genome). If some contigs were in fact closely physically linked on chromosomes, this could lead to larger LD statistics than expected for unlinked loci.

## Discussion

We presented estimators for a range of summary statistics of linkage disequilibrium, including Hill and Robertson’s *D*^2^, *Dz*, and *π*_2_, that account for both phasing issues and finite samples. We were unable to obtain an unbiased estimate of *r*^2^.

Computing estimates and expectations of ratios is challenging, and sometimes intractable. One commonly used approach to estimate *r*^2^ is to first compute 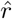 via an EM algorithm (Excoffier and Slatkin, 1995) or genotype covariances (Rogers and Huff, 2009), and then square the result. While we can compute unbiased estimators for *r* from either phased or unphased data, this approach gives inflated estimates of *r*^2^ because it does not properly account for the variance in 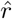. In general, the expectation of a function of a random variable is not equal to the function of its expectation:

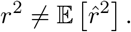

For large enough sample sizes, this error will be practically negligible, but for small to moderate sample sizes, the estimates will be upwardly biased, sometimes drastically (Figures 1 and 2).

An alternative approach is to estimate *D*^2^ and *π*_2_ and compute their ratio for each pair of loci. Given unbiased estimates of the numerator 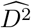 and denominator 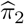, the ratio 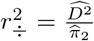 performs favorably to the Rogers-Huff approach (Figure 1C-D) and displays the appropriate decay behavior in the large recombination limit (Figure 2). It is still a biased estimator for *r*^2^, however, since

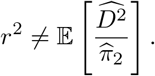

Even if we were given an adequate estimator for *r*^2^, obtaining theoretical predictions for its value is very challenging (McVean, 2002; Song and Song, 2007; Rogers, 2014).

One approach to handle the finite sample bias is to work directly with the finite-sample correlation, i.e., the expected *r*^2^ due to both population-level LD and LD induced by sampling with sample size *n*. This may be estimated by first solving for the expected two locus sampling distribution for a given sample size (Hudson, 2001; Kamm et al., 2016; Ragsdale and Gutenkunst, 2017), and then using this distribution to compute 𝔼[*r*^2^] for that sample size (Spence and Song, 2019). It can also be estimated directly by simulation (e.g., Gutenkunst et al. (2009)). This approach allows for a fair comparison between model expectations and *r*^2^ as computed by the Rogers and Huff (2009) estimator. However, because finite-sample bias dominates signal for all but the shortest recombination distances, comparisons using biased statistics miss relevant patterns of linkage disequilibrium.

Working directly with 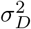, for which we have both unbiased estimators and theoretical predictions, avoids a number of complications. Among caveats for the present approach, we find that the unbiased estimators are analytically cumbersome. For example, expanding 𝔼[*D*^2^] as a monomial series in genotype frequencies results in nearly 100 terms. The algebra is straightforward, but writing the estimator down by hand would be a tedious exercise, and we used symbolic computation to simplify terms and avoid algebraic mistakes. This might explain why such estimators were not proposed for higher orders than *D* in the foundational work of LD estimation in 1970s and 80s. Deriving and computing estimators poses no problem for an efficiently written computer program that operates on observed genotype counts.

For very large sample sizes, however, the bias in the Rogers-Huff estimator for *r*^2^ is weak, and it may be preferable to use their more straightforward approach. Additionally, it is worth noting that like many unbiased estimators, both 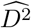 and 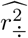 can take values outside the expected range of the corresponding statistics: for a given pair of loci 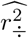 may be slightly negative or greater than one.

Throughout, we assumed populations to be randomly mating. Under inbreeding, there are multiple interpretations of *D* depending on whether we consider the covariance between two randomly drawn haplotypes from the population or consider two haplotypes within the same diploid individual (Cockerham and Weir, 1977). Given the existence of theoretical predictions for two-locus statistics in models with inbreeding, deriving unbiased statistics for this scenario appears a worthwhile goal for future work.

## Methods

### Notation

Variables without decoration represent quantities computed as though we know the true population haplotype frequencies. We use tildes to represent statistics estimated by taking maximum likelihood estimates for allele frequencies from a finite sample: e.g. 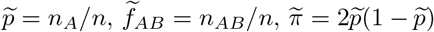, etc. Hats represent unbiased estimates of quantities: e.g. 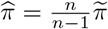. *f*’s denote haplotype frequencies in the population, while *g*’s denote genotype frequencies (Table 2).

**Table 2:** Expected genotype frequencies under random mating. For a given pair or loci, the nine possible two-locus genotypes have frequencies which sum to 1. We assume random mating, so marginal genotype frequencies follow expected Hardy-Weinberg proportions.

|  | *BB* | *Bb* | *bb* | Marginal freq. |
| --- | --- | --- | --- | --- |
| *AA* | $g_1$ | $g_2$ | $g_3$ | $p^2$ |
| *Aa* | $g_4$ | $g_5$ | $g_6$ | $2p(1-p)$ |
| *aa* | $g_7$ | $g_8$ | $g_9$ | $(1-p)^2$ |
| Marginal freq. | $q^2$ | $2q(1-q)$ | $(1-q)^2$ | 1 |

### Estimating statistics from phased data

Suppose that we observe haplotype counts (*n*_*AB*_, *n*_*Ab*_, *n*_*aB*_, *n*_*ab*_), with Σ*n*_*j*_ = *n*, for a given pair of loci. Estimating LD in this case is straightforward. An unbiased estimator for *D* is

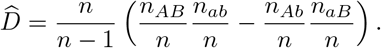

We interpret *D* = *f*_*AB*_*f*_*ab*_ − *f*_*Ab*_*f*_*aB*_ as the probability of drawing two chromosomes from the population and observing haplotype *AB* in the first sample and *ab* in the second, minus the probability of observing *Ab* followed by *aB*. This intuition leads us to the same estimator 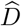:

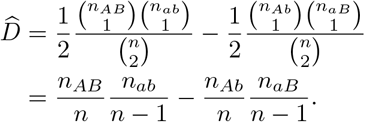

In this same way we can find an unbiased estimator for any two-locus statistic that can be expressed as a polynomial in haplotype frequencies. For example, the variance of *D* is

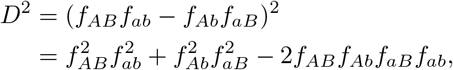

with each term being interpreted as the probability of sampling the given ordered haplotype configuration in a sample of size four (Strobeck and Morgan, 1978; Hudson, 1985). An unbiased estimator for *D*^2^ is then

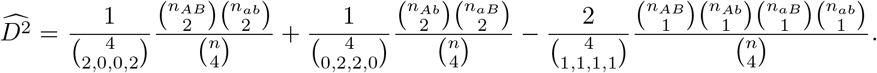

The multinomial factors in front of each term account for the number of distinct orderings of the sampled haplotypes. We similarly find unbiased estimators for the other terms in the Hill-Robertson system, *D*(1 − 2*p*)(1 − 2*q*) and *p*(1 − *p*)*q*(1 − *q*) (shown in the Appendix), or any other statistic that we compute from haplotype frequencies.

### Estimating statistics from unphased data

Estimating two-locus statistics from genotype data requires a bit more work because the underlying haplotypes are ambiguous in a double heterozygote, *AaBb*. Our first step is to derive expressions for *D*, *p*, and *q* in terms of the population genotype frequencies (*g*_1_, …, *g*_9_). We will then use these expressions to derive unbiased estimates in terms of the finite population sample genotype counts (*n*_1_, …, *n*_9_). Expressions for *p* and *q* in terms of genotype frequencies can be read directly from Table 2: 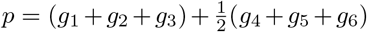, and 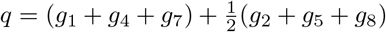. To obtain an estimate for *D* = *f*_*AB*_*f*_*ab*_ − *f*_*Ab*_*f*_*aB*_, we would like to have expressions for haplotype frequencies such as *f*_*AB*_ in terms of the *g*_*i*_.

We can write a naive estimate for *f*_*AB*_ by reading from Table 2 and simply assuming that the double heterozygote genotype *g*_5_ = 2*f*_*AB*_*f*_*ab*_ + 2*f*_*AB*_*f*_*ab*_ had equal probability of the two possible phasing configurations:

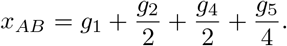

The correct expression for *f*_*AB*_ would replace 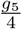 by the probability of the correct haplotype configuration, *f*_*AB*_*f*_*ab*_. This probability can be expressed as 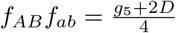, so that

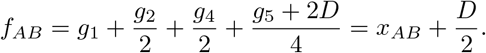

We can obtain similar expressions for all the *f*_··_ and substitute in the expression for *D* to write

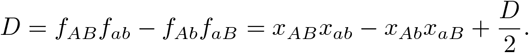

Rearranging provides an estimate of *D* in terms of naive frequency estimates that depend only on genotypes:

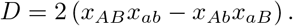

This expression for *D* is equal to Burrows’ “composite” covariance measure of LD,

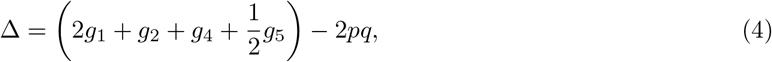

as given in Weir (1979) and Weir (1996), page 126.

Given this expression for D, as well as *p* = *x*_*AB*_ + *x*_*Ab*_ and *q* = *x*_*AB*_ + *x*_*aB*_, we can express higher-order moments as function of genotype frequencies. The Hill-Robertson statistics can be written as polynomials in the naive estimates

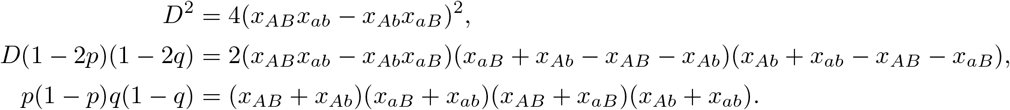

The next step is to obtain estimates from finite samples. Any statistic *S* written as a polynomial in (*x*_*AB*_, *x*_*Ab*_, *x*_*aB*_, *x*_*ab*_) can be expanded as a monomial series in genotype frequencies *g*_*j*_, *j* = 1, …, 9:

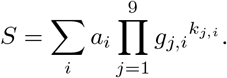

Each term of the form 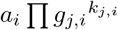 can be interpreted as the probability of drawing *k* = Σ*k*_*j*_ diploid samples, and observing the ordered configuration of *k*_1_ of type *g*_1_, *k*_2_ of type *g*_2_, and so on. Then, from a diploid sample size of *n* ≥ *k*, this term has the unbiased estimator

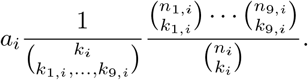

Summing over all terms gives us an unbiased estimator for *S*:

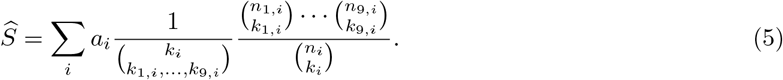

We can use this approach to derive an unbiased estimator for *D*,

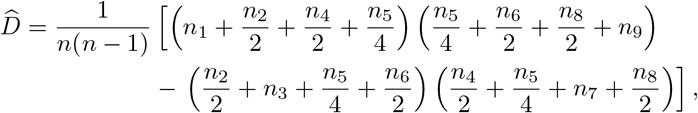

which simplifies to the known Burrows (1979) estimator,

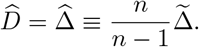

For statistics of higher order than *D*, such as those in the Hill-Robertson system, expanding these statistics often involves a large number of terms. In practice, we use symbolic computation software to compute our estimators. In some cases the estimators simplify into compact expressions, although in other cases they may remain expansive. However, even when there are many terms, the sums do not consist of large terms of alternating sign, and so computation is stable. Mathematica notebooks are provided as supplementary material.

### Simulations of unlinked loci

We used `fwdpy11` (version 0.4.2) (Thornton, 2014) to simulate data with multiple chromosomes and variable population sizes, sample sizes, selfing probabilities, and mutation and recombination rates. To simulate *m* chromosomes each of length *L* base pairs with recombination rate *c* per base pair, we defined *m* segments of “genomic length” 1 with total recombination rate *Lc*, separated with a binomial point probability of 0.5. The total mutation rate was then *mLu*, where *u* is the per-base mutation rate. `fwdpy11` allows the user to define any selfing probability between 0 and 1.

For a given sample size of *n* diploids, we sampled from the *N*_*e*_ simulated individuals without replacement and assumed data from diploids was unphased. To compute 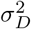 and *σ*_*Dz*_, we constructed genotype arrays and used the `Parsing` features of `moments.LD` (Ragsdale and Gravel, 2019), which makes use of `scikit-allel` (version 1.2.0) (Miles and Harding, 2016), to compute statistics between each pair of chromosomes. We also output the same data in the genepop format, the required input format for `NeEstimator` (version 2.1) (Do et al., 2014), to compute 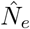 using Equations 1 and 2. For comparisons with Do et al. (2014), we considered minor allele frequency cutoffs of 0.1 and 0.05, as well as using all SNPs.

### Island Fox data and analysis

#### Data

We reanalyzed data for six Channel Island fox populations studied by Funk et al. (2016) (with data deposited at https://datadryad.org/resource/doi:10.5061/dryad.2kn1v). In short, Funk et al. (2016) used Restriction-site Associated DNA sequencing to generate SNP data for between 18 and 46 individuals per population. No reference genome for the island foxes was available at the time, so they generated reference contigs from eight high coverage individuals to map reads from the remaining 192 sequenced individuals. They excluded loci that were called in fewer than half of all individuals and individuals with genotypes for less than half of all loci, and kept only SNPs with minor allele frequency greater than 0.1. They also reported only a single SNP per contig, keeping the first SNP for each contig if more than one SNP were observed.

#### Computing statistics and *N*_*e*_

The sequencing and filtering procedure from Funk et al. (2016) resulted in 5,293 SNPs, which were made available at the above URL. We converted the given genepop format to VCF, and used `scikit-allel` (Miles and Harding, 2016) to parse the VCF and our software `moments.LD` to compute two-locus statistics using the approach described in this paper. We computed both 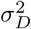 and *σ*_*Dz*_ for each of the six populations. Because contigs were not mapped to a reference genome, we did not know which chromosome each SNP was on. For this analysis, we assumed all SNPs were unlinked.

We computed 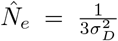 for each population. To compute bootstrapped 95% confidence intervals, we randomly assigned the 5,293 SNPs to 20 groups and computed statistics between all 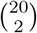 pairs of groups, and then sampled the same number of subset pairs with replacement to compute 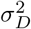 and *σ*_*Dz*_. We repeated this 200 times to estimate the sampling distributions and the 2.5 − 97.5% confidence intervals for each.

### Software

Code to compute two-locus statistics in the Hill-Robertson system is packaged with our software `moments.LD`, a python program that computes expected LD statistics with flexible evolutionary models and performs likelihood-based demographic inference (https://bitbucket.org/simongravel/moments). `moments.LD` also computes LD statistics from genotype data or VCF files using the approach described in this paper, for either phased or unphased data. Code used to compute and simplify unbiased estimators and python scripts to recreate analyses and figures in this manuscript can be found at https://bitbucket.org/aragsdale/estimateld.

## Supporting information

Supporting Mathematica notebook

## Acknowledgements

We thank Lounès Chikhi, Mandy Yao, Alex Diaz-Papkovich, Rosie Sun, and Chris Gignoux for useful discussions. We are grateful to Kevin Thornton for providing helpful guidance in simulating data using `fwdpy11`. This research was undertaken, in part, thanks to funding from the Canada Research Chairs program, the NSERC discovery grant, and CIHR MOP-136855.

## Appendix

## Unbiased estimators from haplotype data

In Methods, we showed how to estimate *D*^2^ from phased data. We can similarly estimate the Hill-Robertson statistics *Dz* = *D*(1 − 2*p*)(1 − 2*q*) and *π*_2_ = *p*(1 − *p*)*q*(1 − *q*):

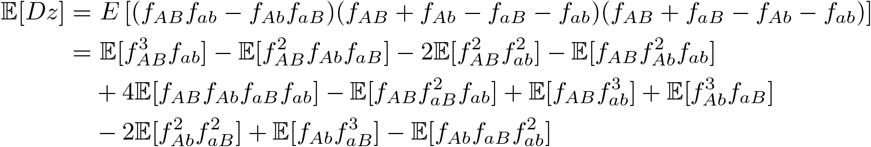

and

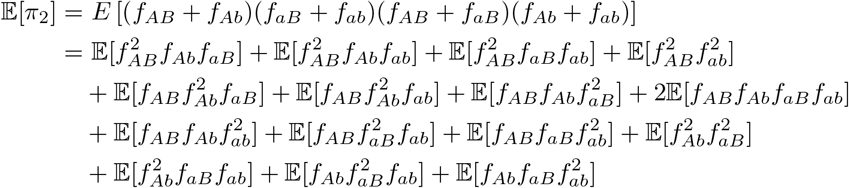

Unbiased estimators for both of these statistics are

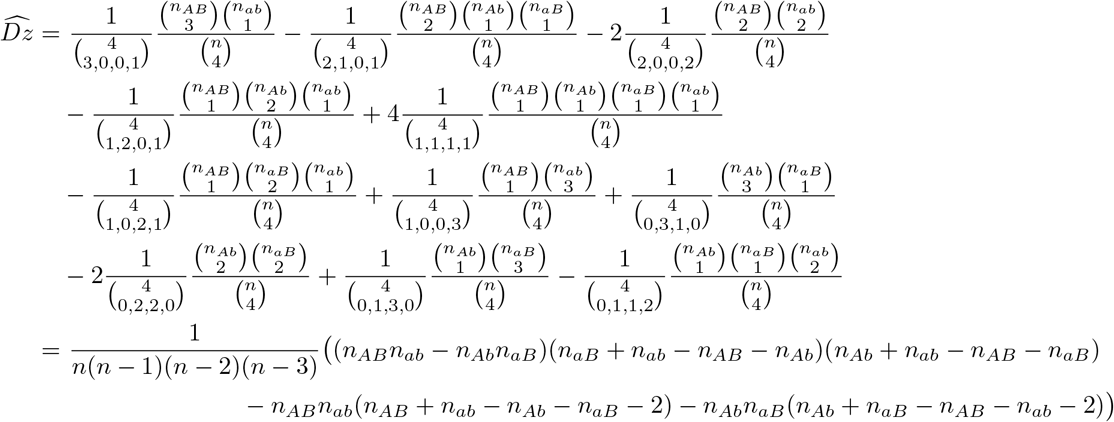

and

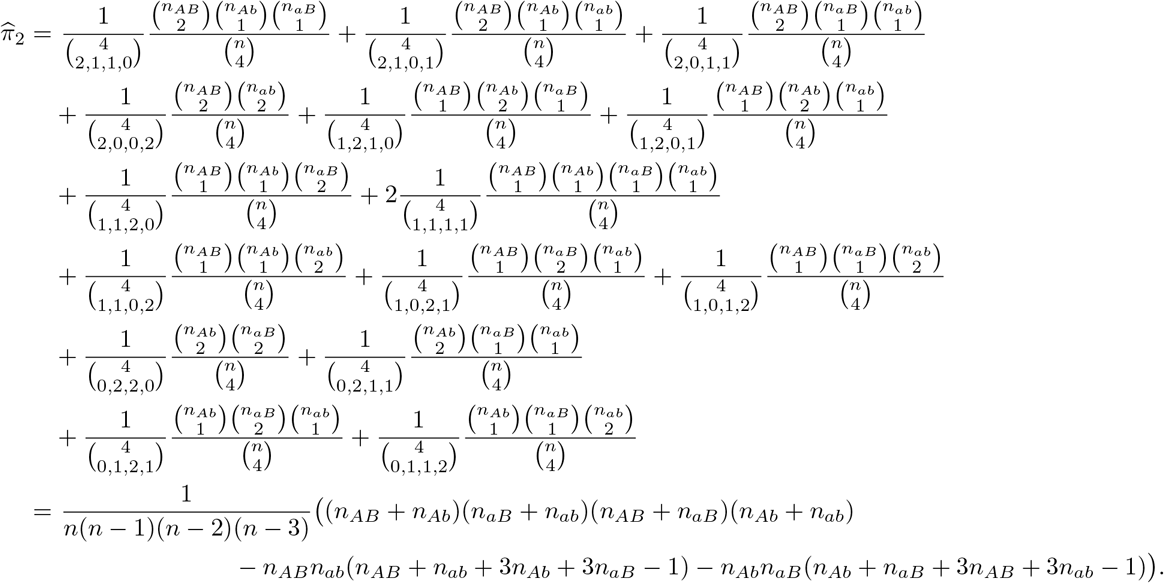

## Multiple populations

We recently presented an extension to the Hill-Robertson system for *D*^2^ to compute expected two-locus statistics across multiple populations, which can be computed rapidly for many related populations with complex demography (Ragsdale and Gravel, 2019). The natural multi-population extension to the Hill-Robertson system (*D*^2^, *Dz*, *π*_2_) is on the basis,

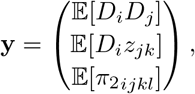

where 1 ≤ *i*, *j*, *k*, *l* ≤ *P* index populations, *z*_*jk*_ = (1 − 2*p*_*j*_)(1 − 2*q*_*k*_), and

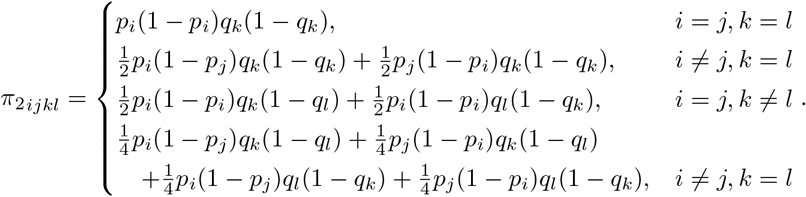

In order to perform inference using these statistics, we estimate them from data and then compare them to their computed expectations. We therefore seek unbiased estimators for each of these statistics from pairs of variable loci across the genome, for either phased or unphased data.

Our approach here is directly analogous to that for computing unbiased estimates of single-population statistics given above, with an additional index over each population. For phased data, we suppose we sample *n*_*i*_ haplotypes from population *i* (1 ≤ *i* ≤ *P*), and from each population we observed haplotype counts (*n*_*i*1_, *n*_*i*2_, *n*_*i*3_, *n*_*i*4_), Σ_*j*_ *n*_*ij*_ = *n*_*i*_. For any statistic *S* as a function of *f*_*ij*_’s, we expand its expectation into a series of monomials, with each term taking the form

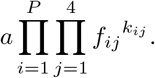

Each term has the interpretation as the probability that we sample Σ_*j*_ *k*_*ij*_ haplotypes from each population, and observe the ordered configuration of *k*_*ij*_ of type *j* in population *i*.

From our sampled haplotypes, an unbiased estimator for each term is

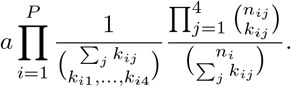

We then sum over each term to obtain our unbiased estimator 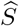.

For example, the covariance of *D* between populations *i* and *j* (*i* ≠ *j*) is given by

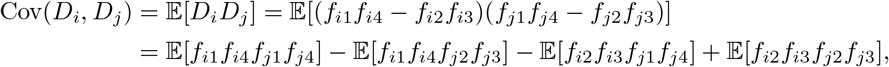

which has the unbiased estimator

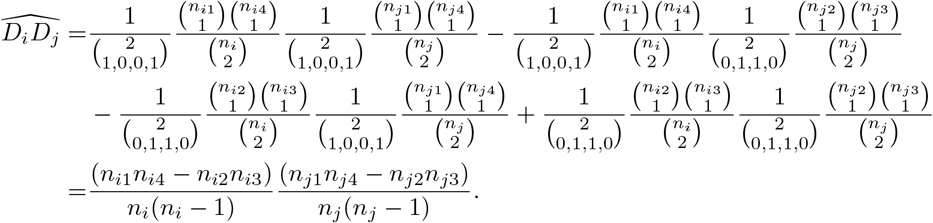

For unphased data, we take the exact same approach, but use genotype frequencies *g*_*ij*_ instead of haplotype frequencies (Table 2), and use the “composite” haplotype frequency estimates in each population *x*_*ij*_, 1 ≤ *j* ≤ 4, as we defined above in the single population case.

Expected statistics on genotype data can be expanded as before as a series of monomials in *g*_*ij*_, with each term taking the form

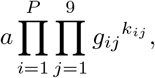

and we again interpret this as the probability of observing a certain genotype configuration in a sample of diploids from each population. Then, just as before, if we sample *n*_*i*_ diploids from each population *i*, with genotype sampling configurations (*n*_*i*1_, …, *n*_*i*9_), an unbiased estimator for any given term is

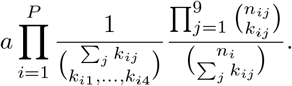

We then sum over terms and simplify to find an unbiased estimator for *S*.

## Expected LD between unlinked loci

In this section, we compute expected 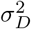 between unlinked loci, first considering the haploid model of Hill and Robertson (1968), followed by a diploid model with random and monogamous mating. These results are similar to predictions from Avery (1978); Weir and Hill (1980) for the related statistic *d*^2^. Our focus on simple mating systems and unlinked loci allows for intuitive derivations for 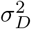 from a genealogical argument.

## Using the Hill-Robertson recursion and hypergeometric sampling

Equations 2 and 3 from Hill and Robertson (1968) give a linear recursion on the statistics 𝔼[*D*^2^], 𝔼[*D*(1 − 2*p*)(1 − 2*q*)], and 𝔼[*p*(1 − *p*)*q*(1 − *q*)], which can be analytically solved at equilibrium (with an influx of mutation, here under the infinite sites model). In Ragsdale and Gravel (2019), we interpreted these two-locus statistics as sampling probabilities, similar to the identity coefficients considered by Hudson (1985) and McVean (2002). Specifically, each term in 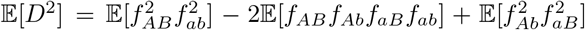 can be thought of as drawing a particular ordered sampling of haplotypes. For example, the first term is the probability of drawing two samples that are each *AB* followed by two samples that are *ab*.

The Hill-Robertson equations describe sampling *with replacement*, but the haplotypes observed in a sequencing experiment are drawn without replacement. If we suppose that samples are chosen randomly from a population of size *N*, we can adjust the expected Hill-Robertson statistics via hypergeometric sampling with known census population size to obtain sampling-*without*-replacement expectations. To do this, we expand each of the Hill-Robertson statistics as expectations over monomial terms in haplotype frequencies, rewrite each term via the hypergeometric distribution with a factor to account for the sampling order, and then simplify again in terms of the Hill-Robertson statistics. For *D*^2^, this is written as

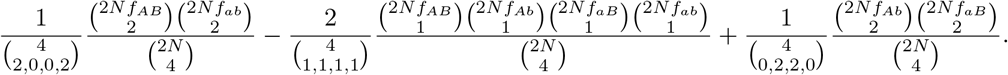

All together, writing

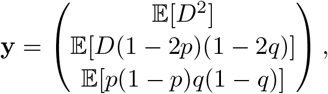

we have

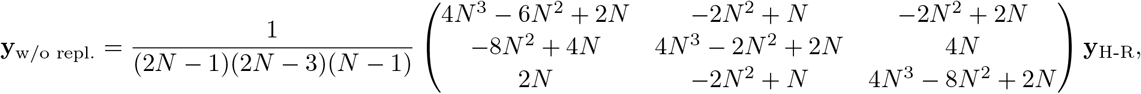

where **y**_H-R_ denote the unadjusted expectations.

From this, we can compute 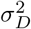 for unlinked loci (*c* = 1/2) under the Hill-Robertson equations assuming sampling without replacement, which gives

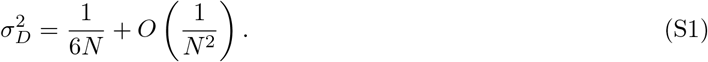

A Mathematica notebook with these calculations is provided as supplementary material.

## Haploid inheritance model

Here, we will approximate 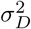 for unlinked loci through an argument based on the probability that two sampled haplotypes are IBD through a recent common ancestor. This approach will rely on a few assumptions. First consider 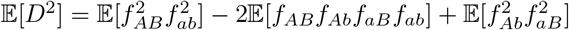, which we illustrate as

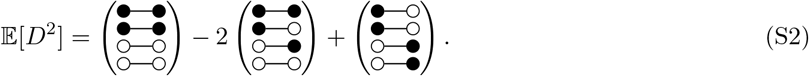

Again, the terms can be thought of as ordered sampling probabilities within a sample (without replacement) of four haplotypes from the population.

If none of the four sampled lineages share a recent common ancestor, we can assume that recombination will have split each lineage before any coalescent event so that the left and right loci coalesce independently. In this case, each term is equally probable, and 𝔼[*D*^2^] = 0.

Alternatively, two of the sampled lineages may share a recent common ancestor (we neglect the case of more than two lineages sharing very recent common ancestors). If two lineages share a recent common ancestor, the sample is expected to contribute nonzero *D*^2^ only if both loci were inherited identical-by-decent (IBD) from the recently shared common ancestor. In this case, we enumerate all the ways to obtain a given 4-haplotype sampling configuration from three haplotypes and an IBD copying event. For example,

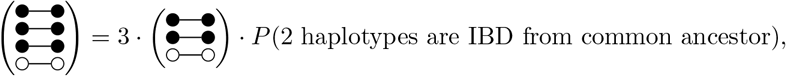

since there are three ways of assigning IBD among the three 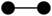 haplotypes in the 4-sample configuration. As another example,

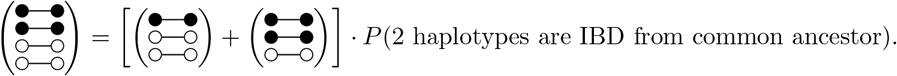

The configuration in the middle term in Equation S2 is not possible if two drawn haplotypes are IBD, so its expectation is zero. We then decompose the remaining terms by conditioning on the probability of inheriting haplotypes IBD through a recent common ancestor and enumerating configurations of possible pairs of shared haplotypes:

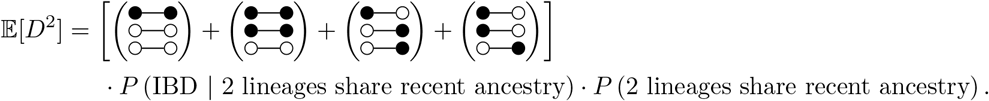

We start with the term in brackets: since the two loci are unlinked, we can assume that population-wide haplotype frequencies are equal to the product of the allele frequencies of the two alleles at the two loci (e.g. 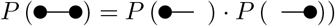). Then, since the three non-IBD haplotypes are inherited independently,

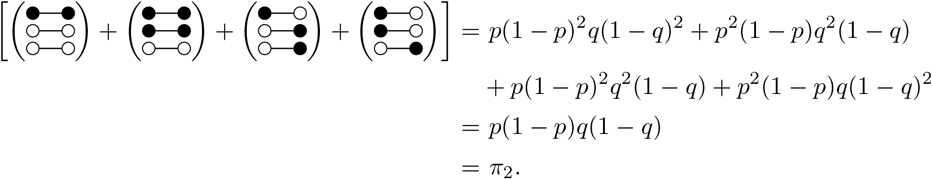

Next, we ask what is the probability that two lineages share a common ancestor some time *t* in the past. In the haploid model, there is no memory of diploid pairings for parents, so that gametes are drawn independently from the population (2*N* total) to form offspring. In the recent past, a gamete has approximately 2^*t*^ genealogical ancestors. The second gamete also has 2^*t*^ ancestors, so each of those ancestors has a 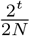 chance of being a shared ancestor to the first gamete. Thus, the probability of sharing a common ancestor *t* generations ago is approximately

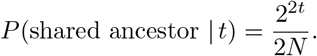

Finally, we compute the probability that both loci were inherited IBD, given that two lineages shared a genealogical ancestor *t* generations ago. In a single generation, there is a 1/4 chance that both loci are inherited from the same parental gamete. Thus, the probability that two lineages are IBD from a common ancestor *t* generations ago is

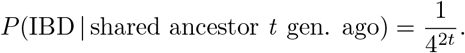

We take this all together, summing over generations, to find

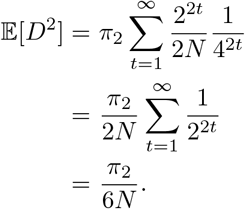

From this,

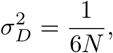

matching to leading order the result of Hill and Robertson (1968) with sampling without replacement (Equation S1).

## Diploid inheritance model

In the Wright-Fisher model, we assume that parents are randomly mating diploid individuals. The difference between this model and the haploid model from the previous section is that while parents are randomly drawn (from *N* total individuals), parents have fixed gametic states. This changes the calculation of the probability of sharing a genealogical ancestor, as we now draw two diploid individuals instead of four independent haploid gametes to generate offspring. It also changes the probability of sharing two unlinked loci IBD.

From the decomposition of 𝔼[*D*^2^] for unlinked loci as above,

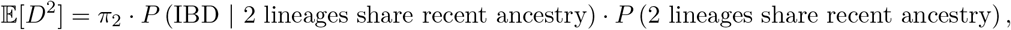

we first compute the probability of sharing a common ancestor *t* generations ago, and then compute the probability of inheriting a pair of unlinked loci IBD from that common ancestor.

First, each sampled haplotype has 2^*t*−1^ ancestors, so any two given samples share a common ancestor in the recent past with probability 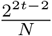. Two offspring of the common ancestor are IBD at two unlinked loci with probability 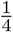. Additional generations reduce the probability of inheriting both loci IBD by 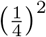, so that the probability of being IBD at the two loci from a common ancestor *t* generations ago is 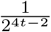.

Combining terms gives us

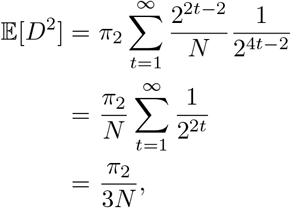

so that

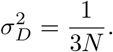

## Monogamy

The model for monogamy is similar to the randomly mating diploid inheritance model in the previous section. Parental pairs are fixed, so siblings always share both parents, and the probability of two randomly drawn haplotypes sharing ancestral (*t*-grand)parents *t* generations ago is 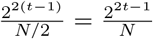. The probability of being IBD at both loci, given that two (*t*-grand)parents were shared *t* generations ago, is 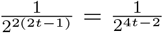. Taken together, we find

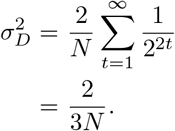

## Expected *σ*_*Dz*_ is approximately zero for unlinked loci

The Hill and Robertson (1968) recursion for 𝔼[*D*^2^] requires also computing the statistic E[*Dz*] = 𝔼[*D*(1 − 2*p*)(1 − 2*q*)]. As discussed in Ragsdale and Gravel (2019), *E*[*Dz*] can be interpreted as *D* weighted by the joint rarity of alleles at the left and right loci. Here, we show that 𝔼[*Dz*], and therefore 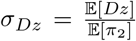, is expected to be zero for unlinked loci.

Following a similar argument to that of 𝔼[*D*^2^] above, we interpret this statistic as a sampling probability as in Equation S2:

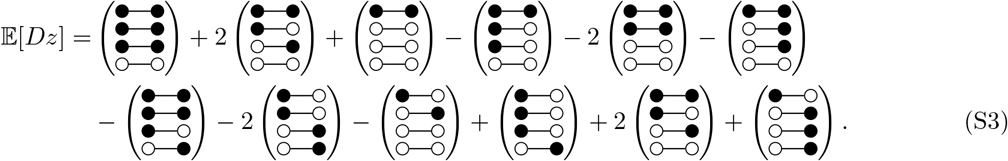

Again, if none of the four sampled lineages share a very recent common ancestor, recombination quickly breaks the association between the two loci, and they are inherited independently. In this case, Equation S3 simplifies to zero, just as it did in Equation S2. Thus, 𝔼[*Dz*] could only be nonzero if multiple lineages share a common ancestor and inherit loci IBD from those share common ancestors.

Consider the case that two lineages share a recent common ancestor from which both loci are inherited IBD. In this case, we can think of all the ways that drawing three lineages and then copying one of them IBD creates the configurations in Equation S3. We assume the three initially drawn lineages do not share a recent common ancestor, so that the left and right loci are inherited independently (again, we ignore higher order terms arising from more than two lineages sharing recent common ancestors). Explicitly writing out this term (and for the moment dropping the probability of pairs of lineages sharing haplotypes IBD), we have

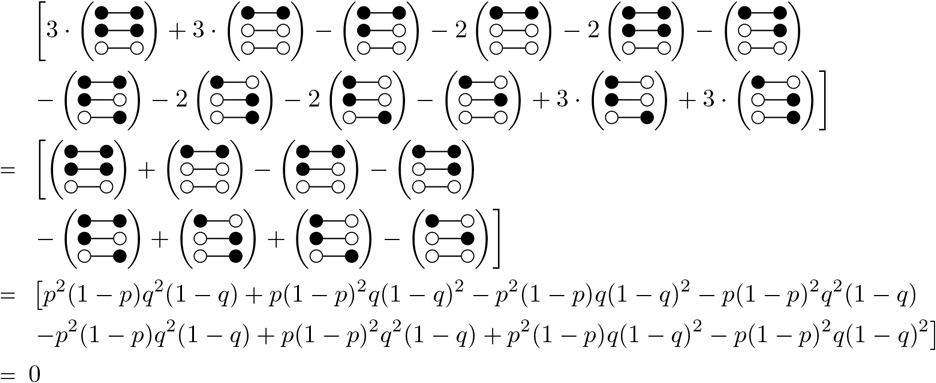

Thus, there is no contribution to 𝔼[*Dz*] from the case of exactly two lineages sharing a recent common ancestor, and so 𝔼[*D*_*z*_] ≈ 0. Considering higher order terms (i.e. more than two lineages sharing very recent common ancestors) leads to 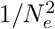 or smaller contributions. This will only result in significantly positive *σ*_*Dz*_ in extremely small population sizes.

## Supplementary figures and tables

**Figure S1:**
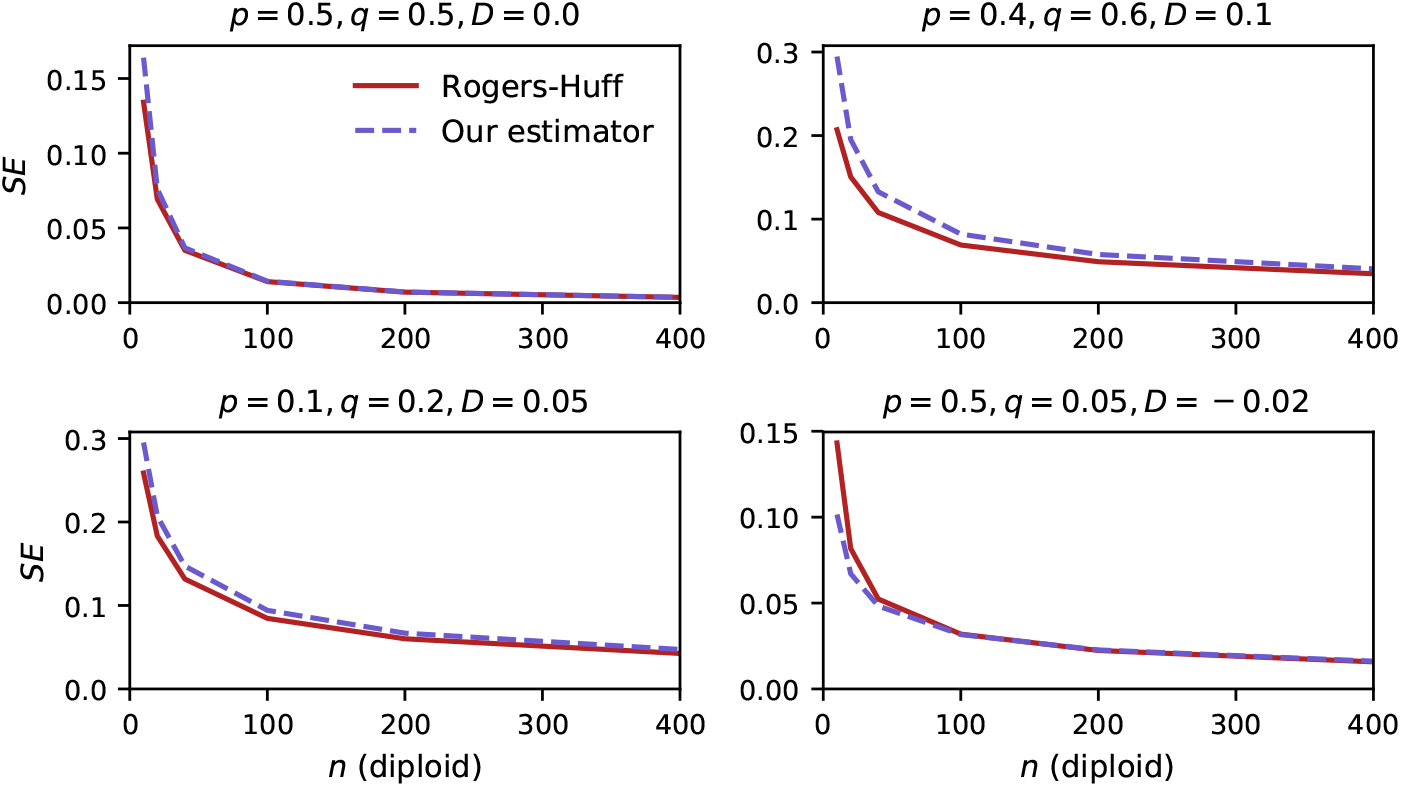
Standard error of *r*^2^ estimators. The Rogers and Huff (2009) estimator for *r*^2^ and our estimator display similar standard errors. However, our estimator is more accurate, especially for small sample sizes (Figure 1).

**Figure S2:**
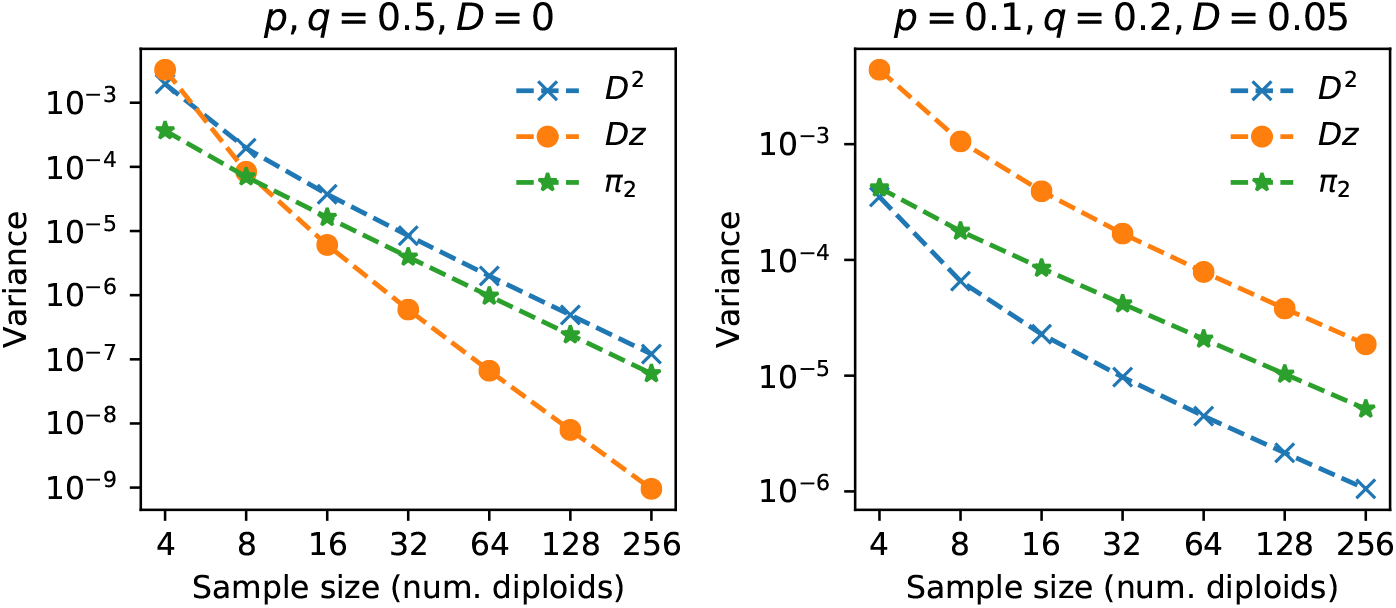
Variance of estimators decays with sample size. Variances decay as 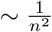 with diploid sample size *n*. These were computed from one million replicates sampled with the given sample size from known haplotype frequencies.

**Figure S3:**
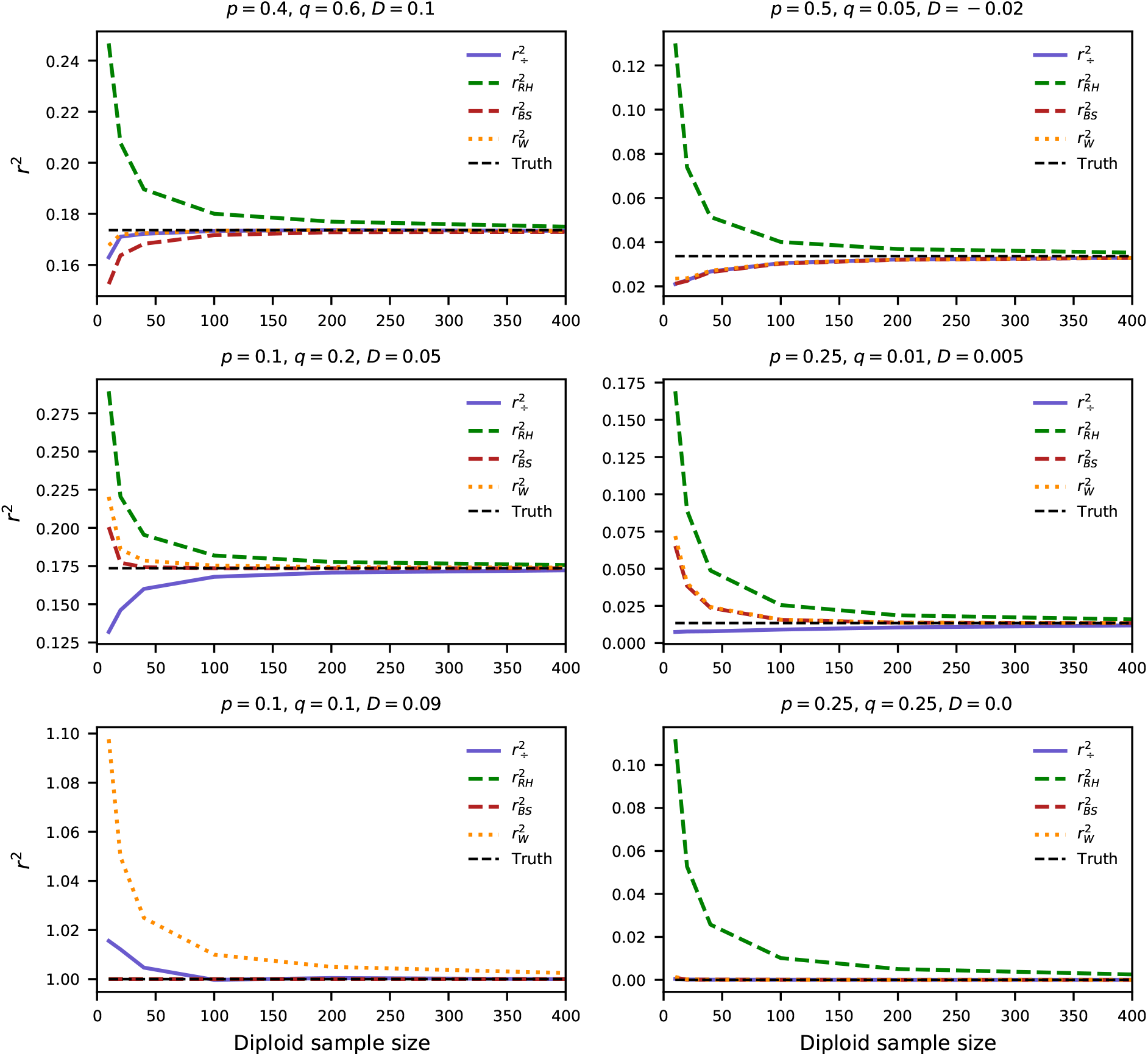
Comparison of estimators for *r*^2^. As in Figure 1C-D, and also including the Bulik-Sullivan et al. (2015) estimator: 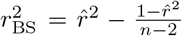, where *n* is the diploid sample size. The Rogers-Huff estimator consistently overestimates *r*^2^ by a large amount, except for the case of perfect LD (lower-left), while the other estimators display variable biases depending on allele frequencies and underly LD.

**Figure S4:**
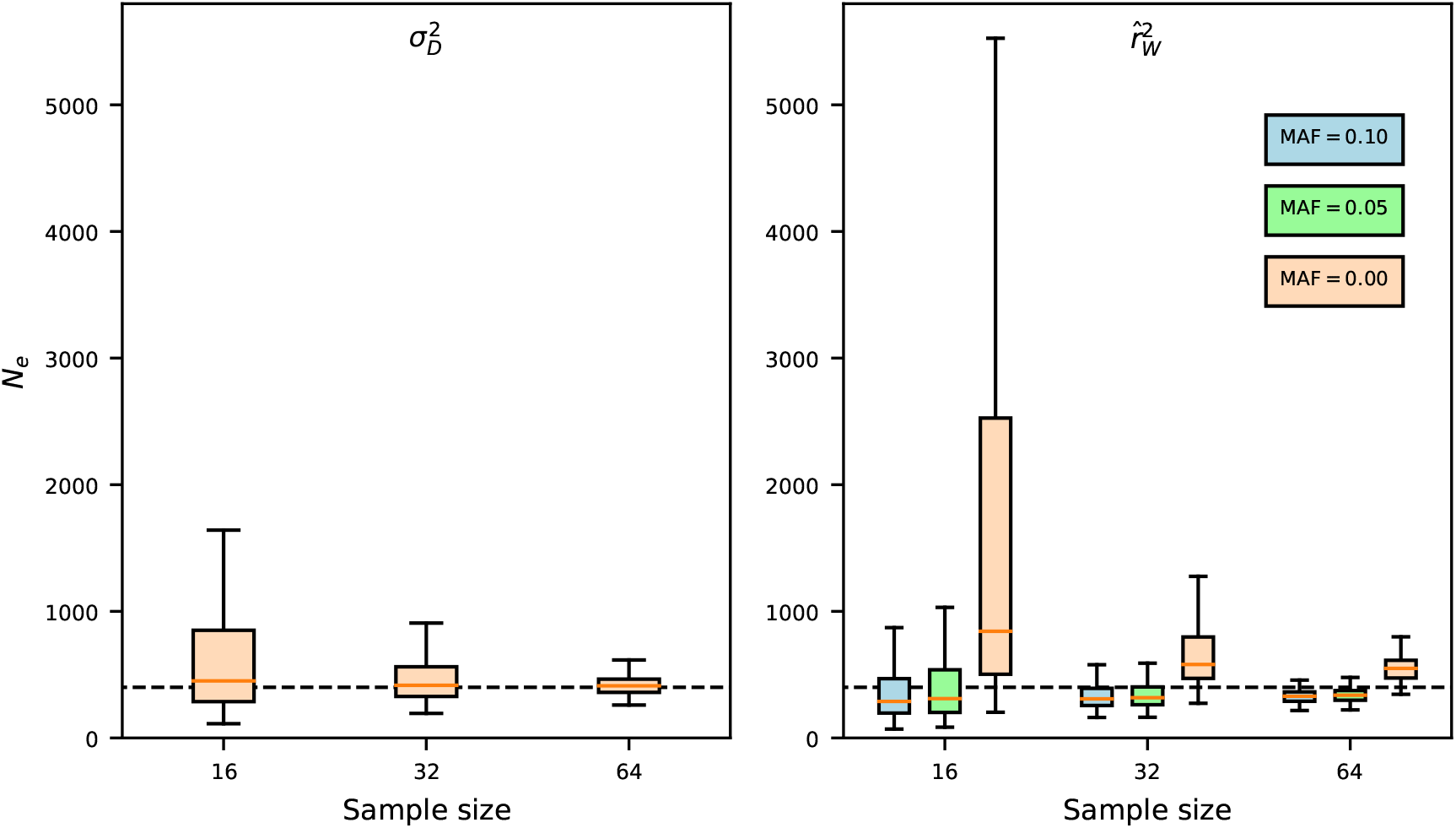
Estimating *N*_*e*_ from unlinked loci, simulated with *N*_*e*_ = 400. We used `fwdpy11` to simulate 10 20Mb chromosomes, with mutation rate *u* = 5 × 10^−8^, recombination rate 2 × 10^−7^, and *N*_*e*_ = 400 (we simulated 200 replicates). From each simulation we sampled 16, 32, and 64 diploids (unphased), and estimated *N*_*e*_ using (left) 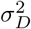 as described here, and (right) `NeEstimator` with minor allele frequency cutoffs 0.1, 0.05, and 0 (i.e., all SNPs). For each method, variance of 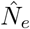 decreased as sample size increased, but `NeEstimator` was biased even for large sample size, with the direction of the bias depending on the chosen MAF.

**Figure S5:**
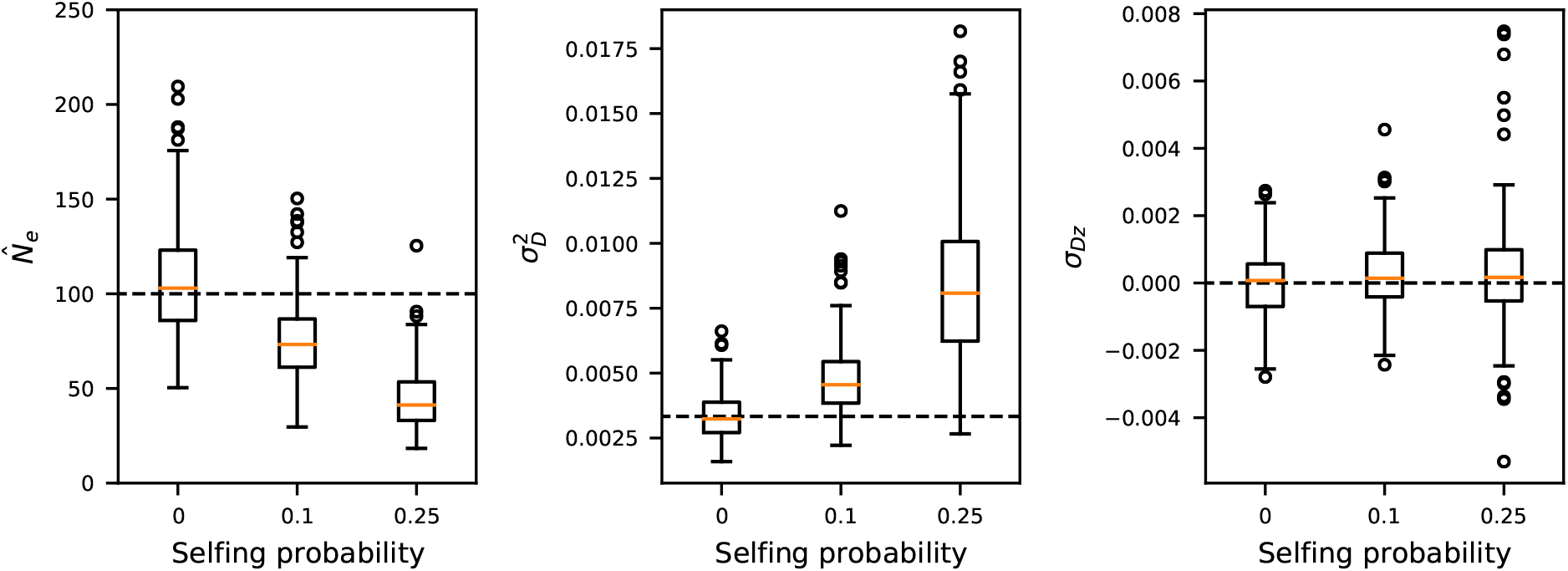
The effect of inbreeding on LD and *N*_*e*_ estimation between unlinked loci. To explore the effects of inbreeding on estimates of 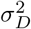 (and thus 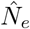) and *σ*_*Dz*_, we used `fwdpy11` to simulate data for two chromosomes, each of length 10^7^ with mutation and recombination rates *u* = *r* = 2×10^−7^ and *N*_*e*_ = 100. To simulate inbreeding, we set the selfing probability to either 0, 0.1, or 0.25. We computed 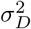 and *σ*_*Dz*_ between all pairs of SNPs from different chromosomes using parsing features in `moments.LD`, and from 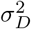 we estimated 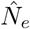. Unsurprisingly, LD as measured by 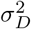 increases with the selfing rate, leading to smaller estimates of 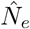. On the other hand, *σ*_*Dz*_ is largely unaffected with higher rates of inbreeding.

**Figure S6:**
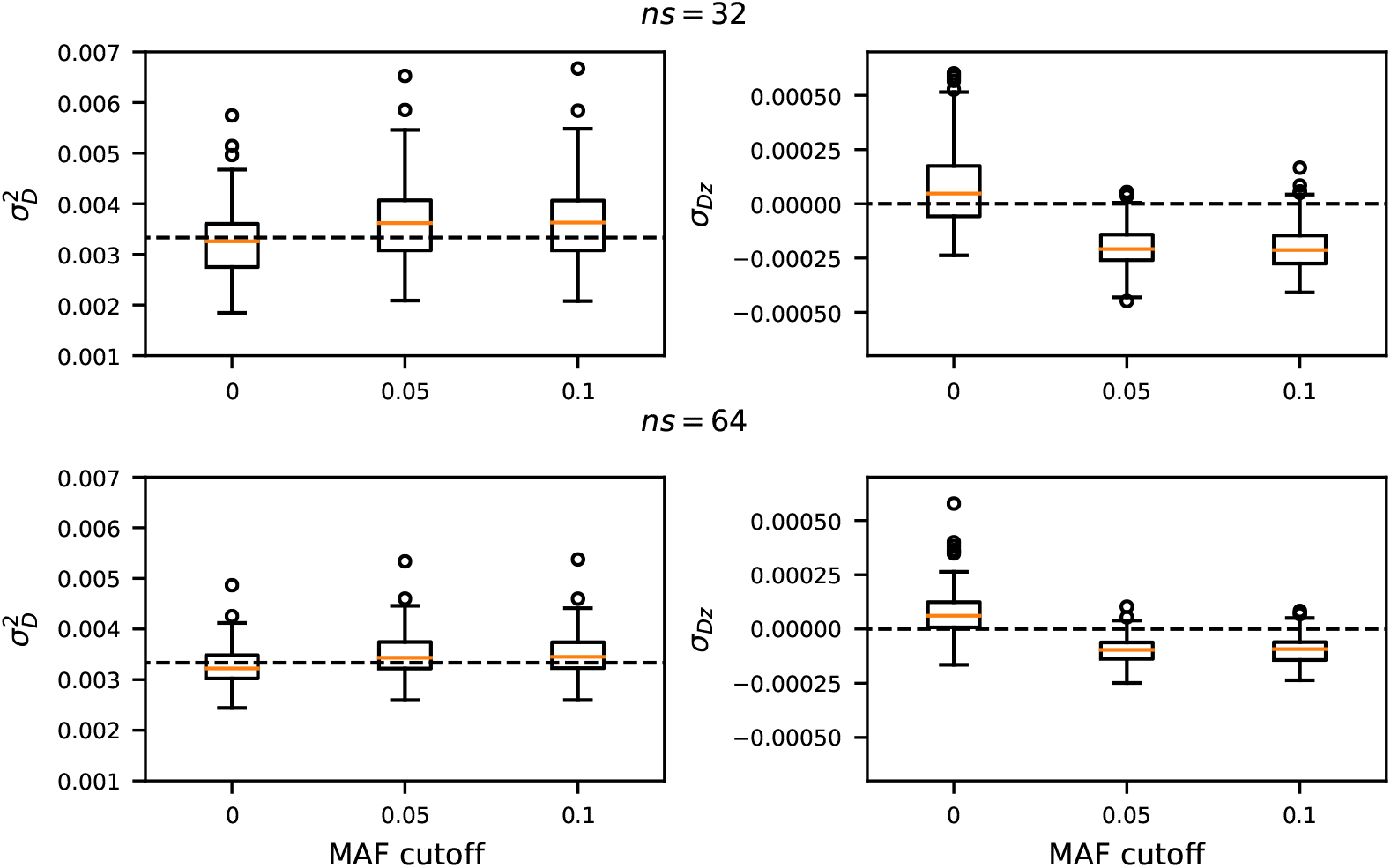
The effect of minor allele cutoff on estimated 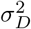 and *σ*_*Dz*_. Using the same simulations as shown in Figure 4, we computed 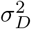 and *σ*_*Dz*_ when filtering data by MAF. With positive MAF cutoff, 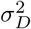 is slightly larger than expectation, while *σ*_*Dz*_ is slightly negative. However, this increase to 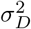 is not large, suggesting that using 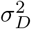 to estimate *N*_*e*_ is less sensitive to filtering by MAF than using *r*^2^ (Figure 4).

**Table S1:**
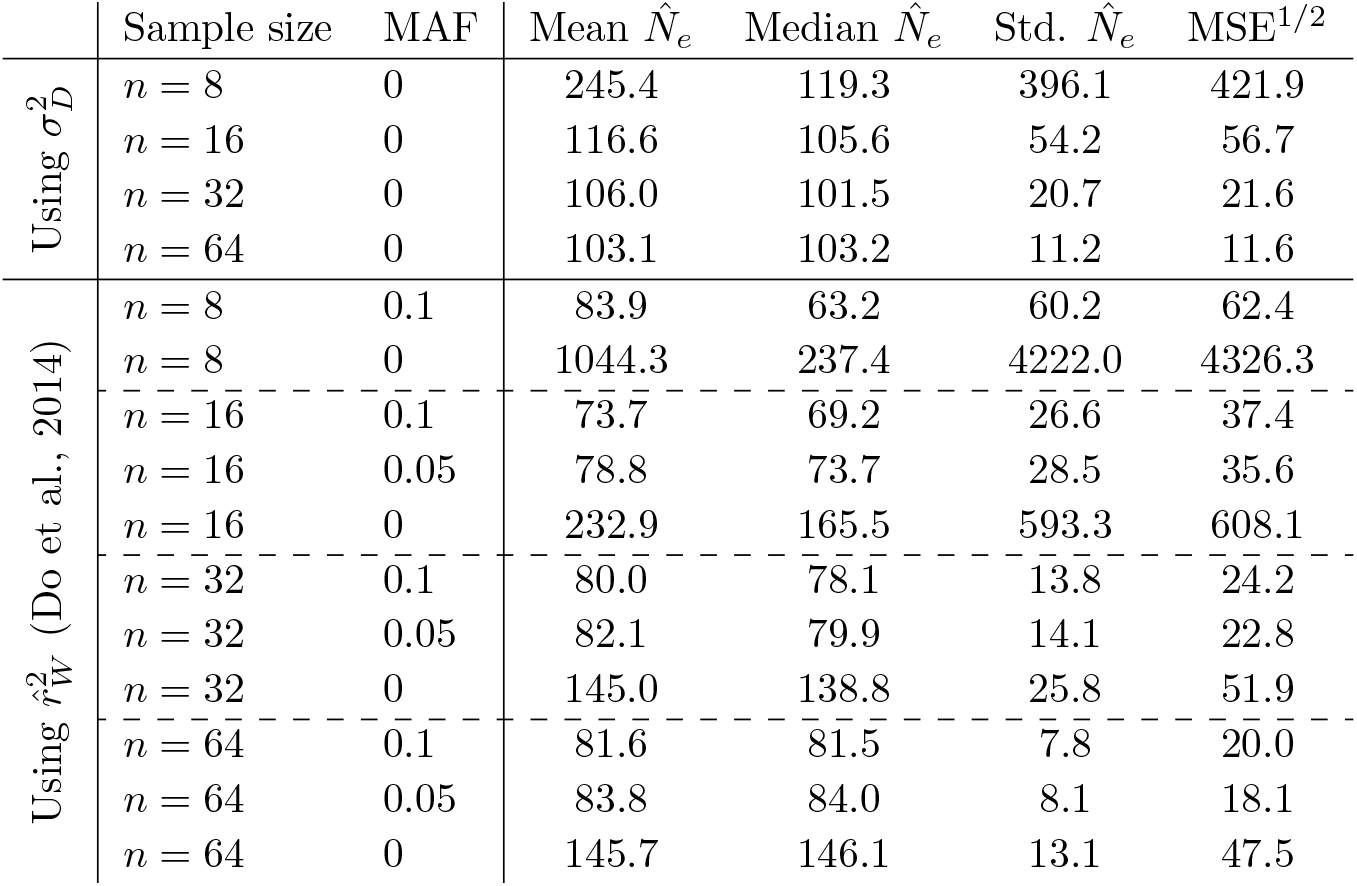
Comparison of 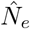 estimation between using 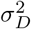 to estimate *N*_*e*_, as proposed in the main text, to (Waples, 2006) estimator for varying minor allele frequency (MAF) cutoffs (Figure 3). 200 simulations were performed using `fwdpy11` (Thornton, 2014) with *N*_*e*_ = 100, 10 chromosomes of length 10 Mb, and recombination and mutation rates *r* = 2 × 10^−7^. In general, using 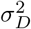 is slightly more variable, but has smaller MSE because it uses an unbiased estimator.

| | Sample size | MAF | Mean $\hat{N}_e$ | Median $\hat{N}_e$ | Std. $\hat{N}_e$ | MSE<sup>1/2</sup> |
| --- | --- | --- | --- | --- | --- | --- |
| Using $\sigma_D^2$ | $n = 8$ | 0 | 245.4 | 119.3 | 396.1 | 421.9 |
| | $n = 16$ | 0 | 116.6 | 105.6 | 54.2 | 56.7 |
| | $n = 32$ | 0 | 106.0 | 101.5 | 20.7 | 21.6 |
| | $n = 64$ | 0 | 103.1 | 103.2 | 11.2 | 11.6 |
| Using $\hat{r}_W^2$ (Do et al., 2014) | $n = 8$ | 0.1 | 83.9 | 63.2 | 60.2 | 62.4 |
| | $n = 8$ | 0 | 1044.3 | 237.4 | 4222.0 | 4326.3 |
| | $n = 16$ | 0.1 | 73.7 | 69.2 | 26.6 | 37.4 |
| | $n = 16$ | 0.05 | 78.8 | 73.7 | 28.5 | 35.6 |
| | $n = 16$ | 0 | 232.9 | 165.5 | 593.3 | 608.1 |
| | $n = 32$ | 0.1 | 80.0 | 78.1 | 13.8 | 24.2 |
| | $n = 32$ | 0.05 | 82.1 | 79.9 | 14.1 | 22.8 |
| | $n = 32$ | 0 | 145.0 | 138.8 | 25.8 | 51.9 |
| | $n = 64$ | 0.1 | 81.6 | 81.5 | 7.8 | 20.0 |
| | $n = 64$ | 0.05 | 83.8 | 84.0 | 8.1 | 18.1 |
| | $n = 64$ | 0 | 145.7 | 146.1 | 13.1 | 47.5 |

**Table S2:** Estimated 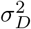 and *σ*_*Dz*_ for island fox populations. Each island fox population exhibited significantly elevated *σ*_*Dz*_, possibly suggesting population substructure or an input of migrants or introduced individuals in recent generations.

| Population | $\sigma_D^2$ | (95% bootstrap CI) | $\sigma_{Dz}$ | (95% bootstrap CI) |
| --- | --- | --- | --- | --- |
| San Miguel I. | 0.0218 | (0.0205 – 0.0231) | 0.0256 | (0.0235 – 0.0284) |
| Santa Rosa I. | 0.0250 | (0.0245 – 0.0256) | 0.00885 | (0.00799 – 0.00963) |
| Santa Cruz I. | 0.0146 | (0.0143 – 0.0149) | 0.00383 | (0.00312 – 0.00454) |
| Santa Catalina I. | 0.00814 | (0.00801 – 0.00824) | 0.00409 | (0.00383 – 0.00437) |
| San Clemente I. | 0.00564 | (0.00489 – 0.00643) | 0.00972 | (0.00802 – 0.0114) |
| San Nicolas I. | 0.0240 | (0.0218 – 0.0258) | 0.00282 | (0.000812 – 0.00494) |

